# A mechanistic, stigmergy model of territory formation in asocial animals: Territorial behavior can dampen disease prevalence but increase persistence

**DOI:** 10.1101/796045

**Authors:** Lauren A. White, Sue VandeWoude, Meggan E. Craft

## Abstract

Mechanistic portrayals of how animals form and maintain territories remain rare, and as a discipline, collective movement ecology has tended to focus on synergistic (e.g., migration, shoaling) rather than agonistic or territorial interactions. Here we ask how dynamic territory formation and maintenance might contribute to disease dynamics in an asocial territorial animal for an indirectly transmitted pathogen. We developed a mechanistic individual-based model where stigmergy—the deposition of signals into the environment (e.g., scent marking, scraping)—dictates not only local movement choices and long-term territory formation, but also the risk of pathogen transmission. Based on a variable importance analysis, the length of the infectious period was the single most important variable in predicting outbreak success, maximum prevalence, and outbreak duration. Population size and rate of pathogen decay were also key predictors. We found that territoriality best reduced maximum prevalence in conditions where we would otherwise expect outbreaks to be most successful: slower recovery rates (i.e., longer infectious periods) and higher conspecific densities. However, at high enough densities, outbreak duration became considerably more variable. Our findings therefore support a limited version of the “territoriality benefits” hypothesis—where reduced home range overlap leads to reduced opportunities for pathogen transmission, but with the caveat that reduction in outbreak severity may increase the likelihood of pathogen persistence. For longer infectious periods and higher host densities, key trade-offs emerged between the strength of pathogen load, strength of the stigmergy cue, and the rate at which those two quantities decayed; this finding raises interesting questions about the evolutionary nature of these competing processes and the role of possible feedbacks between parasitism and territoriality. This work also highlights the importance of considering social cues as part of the movement landscape in order to improve our understanding of the consequences of individual behaviors on population level outcomes.

**Author summary:** Making decisions about conservation and disease management relies on our understanding of what allows animal populations to be successful. However, movement ecology, as a field, tends to focus on how animals respond to their abiotic environment. Similarly, disease ecology often focuses on the social behavior of animals without accounting for their individual movement patterns. We developed a simulation model that bridges these two fields by allowing hosts to inform their movement based on the past trajectories of other hosts. As hosts navigate their environment, they leave behind a scent trail while avoiding the scent trails of other individuals. We wanted to know if this means of territory formation could heighten or dampen disease spread when infectious hosts leave pathogens in their wake. We found that territoriality could inhibit disease spread under conditions that we would normally expect pathogens to be most successful: where there are many hosts on the landscape and hosts stay infectious for longer. This work points to how incorporating movement behavior into disease models can provide improved understanding of how diseases spread in wildlife populations; such understanding is particularly important in the face of combatting ongoing and emerging infectious diseases.

## Introduction

In their movement ecology framework, Nathan et al. [1] suggest four motivating questions: (1) why move?; (2) how to move?; (3) when and where to move?; and (4) what are the ecological and evolutionary consequences of moving? Recently, there has been a call for the discipline of movement ecology to better address the fourth component of this framework: the population-level consequences of moving [2]. In particular, researchers have argued for a greater synthesis of movement ecology with biodiversity [3] and disease ecology research [4, 5]. One of the goals of incorporating such detail is to be able to observe the emergence of complex ecological and evolutionary processes that may depend upon individual traits like personality or behavioral phenotypes [6]. Pathogen transmission is one such process that is highly dependent on whether two conspecifics encounter each other within a certain window of time and space.

However, as a discipline, movement ecology has largely focused on individual movement as a function of environmental or social drivers, but rarely both [7]. Collective movement ecology, in particular, has focused on synergistic movements among group members (e.g., shoaling, migration) [8]. In contrast, mechanistic portrayals of how animals form and maintain territories remain rare [9] (but see: [10–12]). Disease ecology faces similar barriers to incorporating contact behavior that explicitly reflects individual movement patterns [4, 5, 13]. Models in disease ecology are often specific to a given-host pathogen system or emphasize the risk of contact rather than ongoing transmission dynamics [5].

These disciplinary trajectories are problematic because wildlife vary in social organization on axes of gregariousness (group living vs. asocial) and territoriality (territorial vs. nonterritorial), and each population structure has its own potential effects on pathogen transmission [14, 15]. In an evolutionary context, parasites are a possible cost of group living, and host gregariousness is hypothesized to correlate with increased parasite prevalence, infection intensity, and parasite species richness [16, 17]; however, this hypothesis has equivocal empirical support [16, 17]. In particular, host movement and territorial behavior may confound the relationship between group size and prevalence of parasitism [15]. A corollary to this idea is that populations with smaller groups or spatially structured populations may be more protected from parasite transmission from external groups [18].

One possible mechanism for the maintenance of territories and spatial structure within populations is stigmergy. Stigmergy describes environmentally mediated feedback where the signals that one individual leaves in its path alter the behavior of its conspecifics, even after the individual has left that location [11]. In social insects, stigmergy helps to explain how individual pheromone trails can shape social organization of colonies [19]. In territorial animals, equivalent cues include marking through urine, scat, or community scrapes [20–23]. For example in puma (*Puma concolor*), males alter their visitation rates to community scrapes depending on the presence or absence of females or male competitors [24]. In a disease context, these non-contact territorial defense strategies (e.g., vocalization, scent marking, scraping) may have evolved to reduce transmission risk between individuals or groups [25, 26]. Social and spatial structure, potentially mediated by such signaling, is one hypothesis for why population thresholds lack strong empirical support in wildlife populations [26–28]. Population thresholds are a key concept in epidemiology and disease ecology and lie at the root of disease control focused on reducing a susceptible population size through culling or vaccination to reduce the likelihood of outbreaks [27].

However, the relationship between population thresholds and indirectly transmitted pathogens remains an open question [27]. Feedbacks between host behavior and parasitism are likely to complicate this relationship further. Hosts have evolved defenses and avoidance behaviors in response to high parasitism risk (e.g., altering of ranging patterns in primates or selective foraging with behavioral avoidance of fecal-contaminated areas in ungulates) [15]. However, have pathogens co-evolved to counteract territorial barriers and exploit social signaling behaviors? Some preliminary evidence suggests that this may occur. For example, wild banded mongooses (*Mungos mungo*) transmit the mycobacterium, *M. mungo*, almost solely through anal and urine secretions, which are key currencies in their social communication system [29]. Similarly, higher rates of raccoon roundworm (*Baylisascaris procyonis*) infection occur at latrine sites compared to individual raccoon sites; this could lead to higher infection rates for susceptible raccoons and intermediate hosts attracted by undigested seeds [30]. While there is preliminary empirical evidence about the potential role of social signaling behaviors in indirect pathogen transmission, we lack a clear understanding of whether stigmergy is a potential mitigator or facilitator of pathogen transmission at a population level.

Here we developed a generalizable mechanistic framework that examines the interplay between indirect pathogen transmission and dynamic territory formation motivated by deposition and response to signals left in the environment by hosts, i.e., stigmergy cues. We scale up the consequences of these individual decisions to simulate movements, interactions, and pathogen spread across a population. We then ask: (1) how do pathogens spread in populations responding to stigmergy stimuli (e.g., scent/territorial marking) compared to populations where individuals move randomly?; and (2) what are the consequences in trade-offs between strength and duration of scent mark vs. pathogen load deposited in the environment? Here we explore the potential role of stigmergy not only in dynamic territory formation [8,10], but as a potential mitigator or facilitator of pathogen transmission in populations.

## Results

### Individual-based stigmergy model with pathogen transmission

We developed a stochastic Susceptible-Infected-Recovered (SIR) model for a closed population (no births, deaths, immigration, or emigration) [31]. This model operates in discrete space and discrete time. At each time step, individuals could move within a Moore neighborhood (8 neighboring cells) or remain within their current cell. Movement was weighted by the simplest case of a movement kernel where weight was inversely proportional to the radial distance from the current location (see Methods for further details).

Hosts navigated the landscape randomly based on this weighted distance kernel, unless an individual encountered a scent marker from another individual during the previous time step (Fig 1, t_0_). At each time step, every individual deposited a scent mark with initial intensity, *η_o_*, at their current location (Fig 1). If an individual was infected, it also deposited pathogens into the environment with intensity, *ĸ_o_*. Pathogen load was cumulative—so if two infected individuals visit the same cell in sequence, the pathogen load in the environment reflected the sum of their two visits. Both scent mark strength and pathogen load in the environment decayed exponentially through time. The current scent mark strength since the time of deposit (*t_d_*) was given by: *η*(*t*) = η_0_e^−*δ*(*t*–*t_d_*)^. Likewise the pathogen load intensity in a cell since the time of deposit (*t_d_*) was given by: *κ*(*t*) = κ_0_e^−*α*(*t–t*_d_)^.

**Fig 1.**
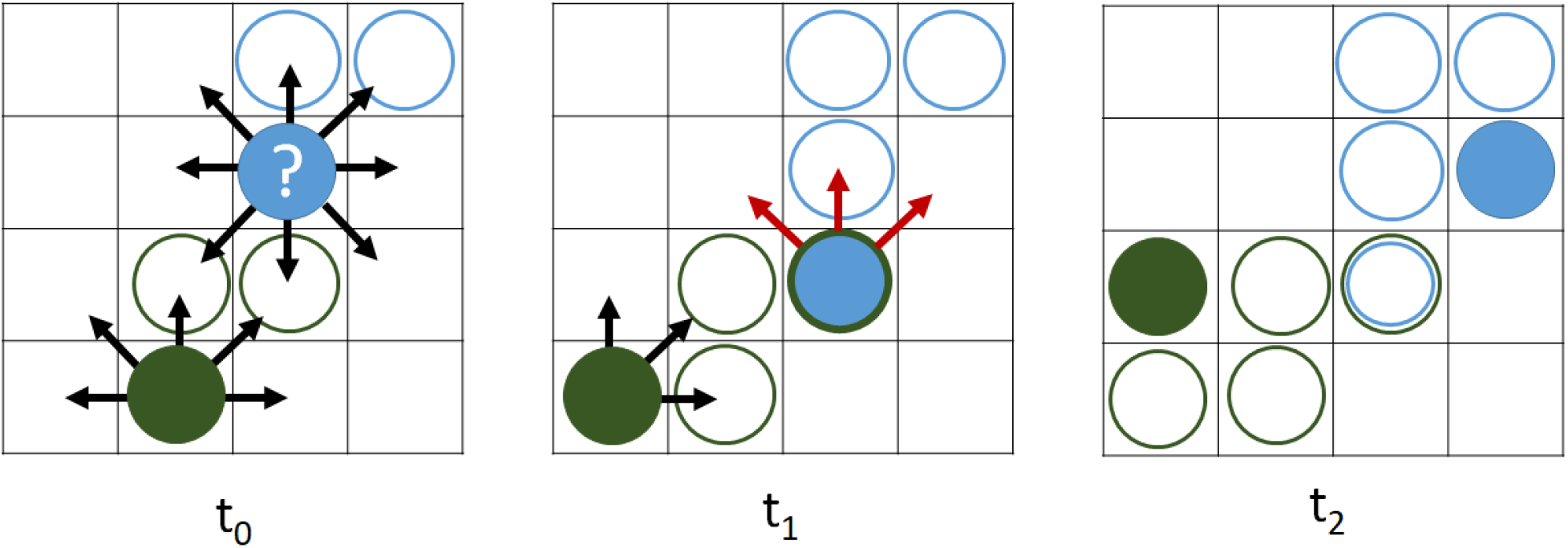
Schematic for individual-based stigmergy model. As hosts (solid circles) walk randomly through space they deposit scent marks (open circles); note the green vs. blue scent marks for each host. If infected, the hosts simultaneously leave pathogens in the environment. **t_0_:** focal individual (blue) can move to one of eight neighboring cells or remain within the current cell. **t_1_**: In the case of encountering a conspecific cue (open green circle), the direction of movement is constrained to 45 degrees on either side of the direction of movement from the previous time step indicated by the directions of the red arrows. **t_2_**: avoiding conspecific stigmergy cues results in dynamic home range formation, but also potential pathogen exposure.

The hosts’ movement responses to these stimuli depended on the strength of the scent load encountered. An individual’s scent exposure was taken as: *min*(1,*Ση_xy_*(*t*)), where *Ση_xy_*(*t*) represents the sum of all active scent load deposited by all hosts in cell location (x, y) on the landscape at time t. To incorporate stochasticity that reflected the strength of the scent cue encountered, a Bernoulli trial determined the subsequent direction of movement. If p < *min*(1, *Ση_xy_*(*t*)), the direction of movement was constrained to 45 degrees on either side of the direction of movement that brought that animal to the current cell (Fig 1, t_1_). For example, a host encountering a foreign scent mark after moving to the bottom middle cell could move to the upper left, upper, or upper right from the current, scent marked cell (Fig 1, t_1_−t_2_). If p > *min*(1, *Ση_xy_*(*t*)), the direction of movement was random, as described by the weighted distance kernel.

We simulated the spread of an indirectly transmitted pathogen, beginning each simulation with a single, infected index case. For susceptible individuals in cells with environmental contamination, the probability of transmission was governed by a binomial distribution where *p* = *min*(1,*Σκ_x,y_*(*t*)), where *Σκ_xy_*(*t*) is the sum of active pathogen load from all previous infected individuals visiting that cell. This type of lattice model of territory formation results in dynamic territories that change through time (Fig 2) [9–11].

**Figure 2.**
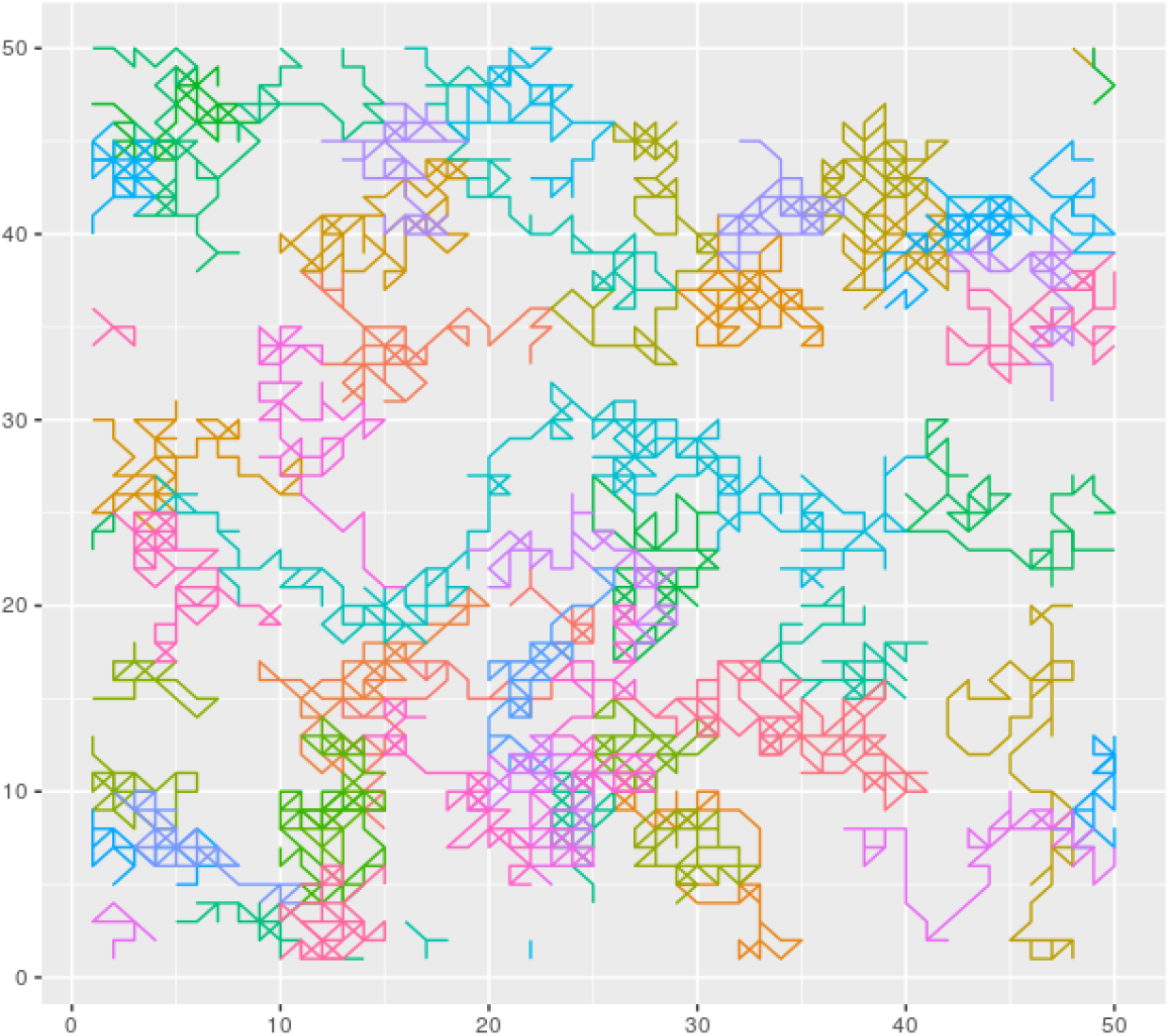
Movement trajectories for simulated hosts. Simulations occurred on a 50 × 50 landscape with wrapped edges (i.e., torus) to avoid boundary effects. Populations were simulated with 50 or 100 individuals. Trajectories shown for a simulation with 50 hosts, each represented by a different color, and a scent cue decay rate of 0.01 per time step for 100-time steps.

Our motivating question was how this dynamic territory formation interacts with indirect pathogen transmission on the landscape. We varied host density (N), recovery rate (*γ*), initial pathogen load(*κ_o_*), pathogen decay rate (*α*), initial scent load (*η_o_*), and scent decay rate (*δ*). We also compared these stigmergy-driven, territorial simulations with their random movement equivalents (*m*). In total, we tested 1,458 parameter sets to with 100 simulations per parameter set (Table 1; see Methods for additional details). For each parameter set, we recorded outbreak success (did the disease spread beyond the initial index case?), maximum prevalence, and outbreak duration (the number of time steps until there were no remaining infectious individuals on the landscape). We also used a random forest variable importance analysis to assess the relative importance of each parameter on these three outcomes.

**Table 1.** Factorial design of 1,458 parameters encompassing host density, recovery rate, pathogen load and decay rate, and scent load and decay rate.

| Parameter | Values tested | Description |
| --- | --- | --- |
| $N$ | 50, 100, 150 hosts | Initial population size |
| $\gamma$ | 0.01, 0.05, 0.10 time<sup>-1</sup> | Recovery rate |
| $m$ | TRUE, FALSE | Stigmergy driven (T) or random movement (F) |
| $\kappa_o$ | 0.1, 1, 10 | Pathogen load: initial strength of pathogen load deposited into the environment |
| $\eta_o$ | 0.1, 1, 10 | Scent load: Initial strength of scent mark deposited into environment |
| $\alpha$ | 0.01, 0.1, 1 | Pathogen decay rate: decay rate of pathogen infectiousness in environment |
| $\delta$ | 0.01, 0.1, 1 | Scent decay rate: decay rate of scent mark strength |

### Recovery rate critical to spread of indirectly transmitted pathogens

The random forest variable importance analysis indicated that recovery rate (*γ*) was the single most important variable in predicting the probability of a successful outbreak, maximum prevalence, and outbreak duration (Fig 3). Population size (N) and decay rate of pathogen infectiousness (*α*) followed as the next most important variables for predicting all three outbreak metrics (Fig 3).

**Fig 3.**
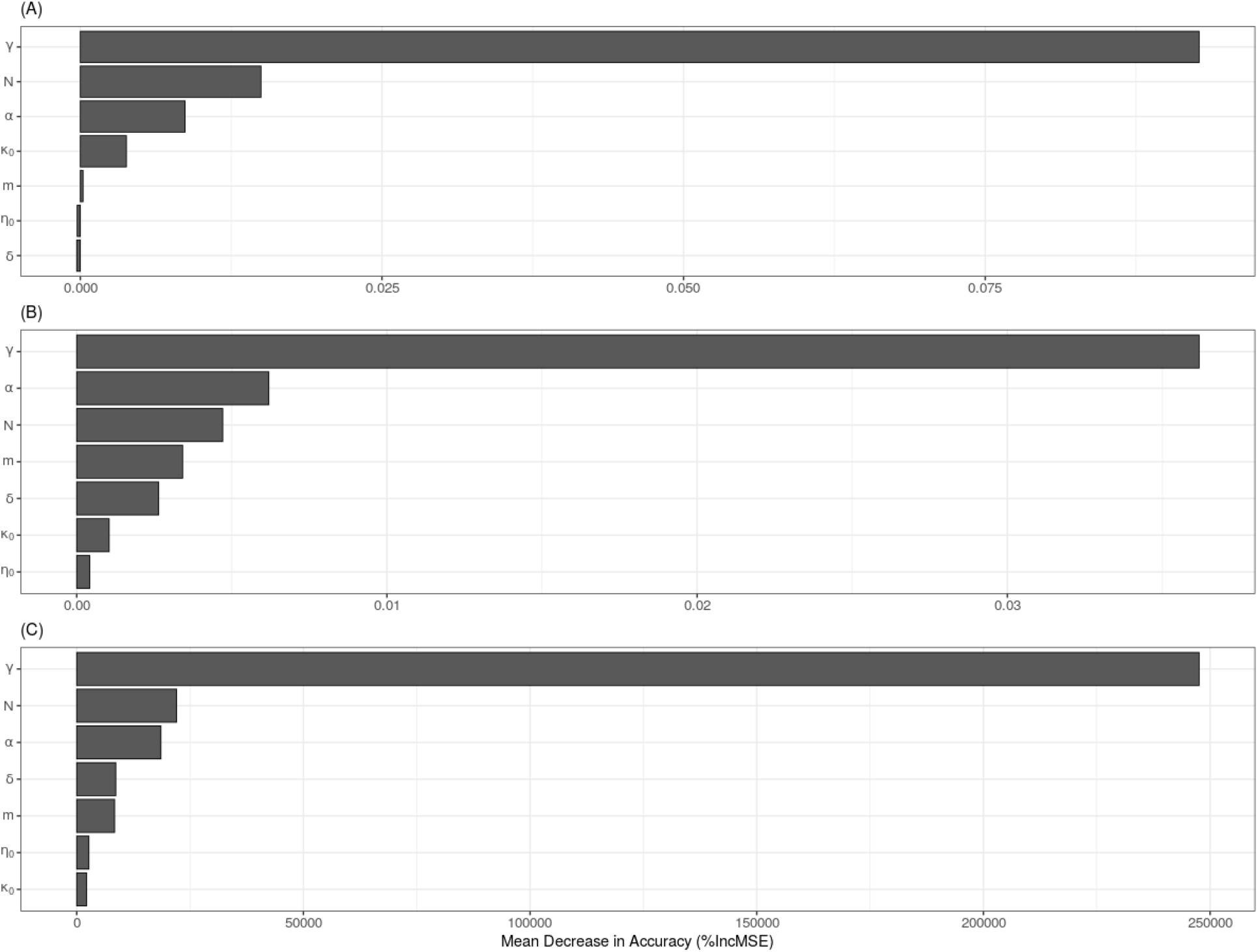
Random forest analysis for: (A) likelihood of a successful outbreak (spreading beyond the index case); (B) maximum prevalence; and (C) outbreak duration. Variable abbreviations: recovery rate (*γ*), population size (*N*), decay rate of pathogen infectiousness (*α*), initial strength of pathogen load deposited into the environment (*κ_o_*), stigmergy-driven vs. random movement (*m*), initial strength of scent mark deposited into environment (*η_o_*), and decay rate of scent mark strength (*δ*).

However, for maximum prevalence specifically, pathogen decay rate (*α*) slightly exceeds population size (N) in variable importance (Fig 3B). Whether or not an outbreak had stigmergy-driven vs. random movement (*m*) had little impact on whether or not an outbreak took place (Fig 3A), but did contribute to determining the maximum prevalence and duration of successful outbreaks (Fig 3A&B). Outbreaks at with faster recovery rates (i.e., 0.1 and 0.05 per time step) had lower maximum prevalence and shorter outbreak durations regardless of whether movement was random or driven by stigmergy cues (Fig 4).

**Fig 4.**
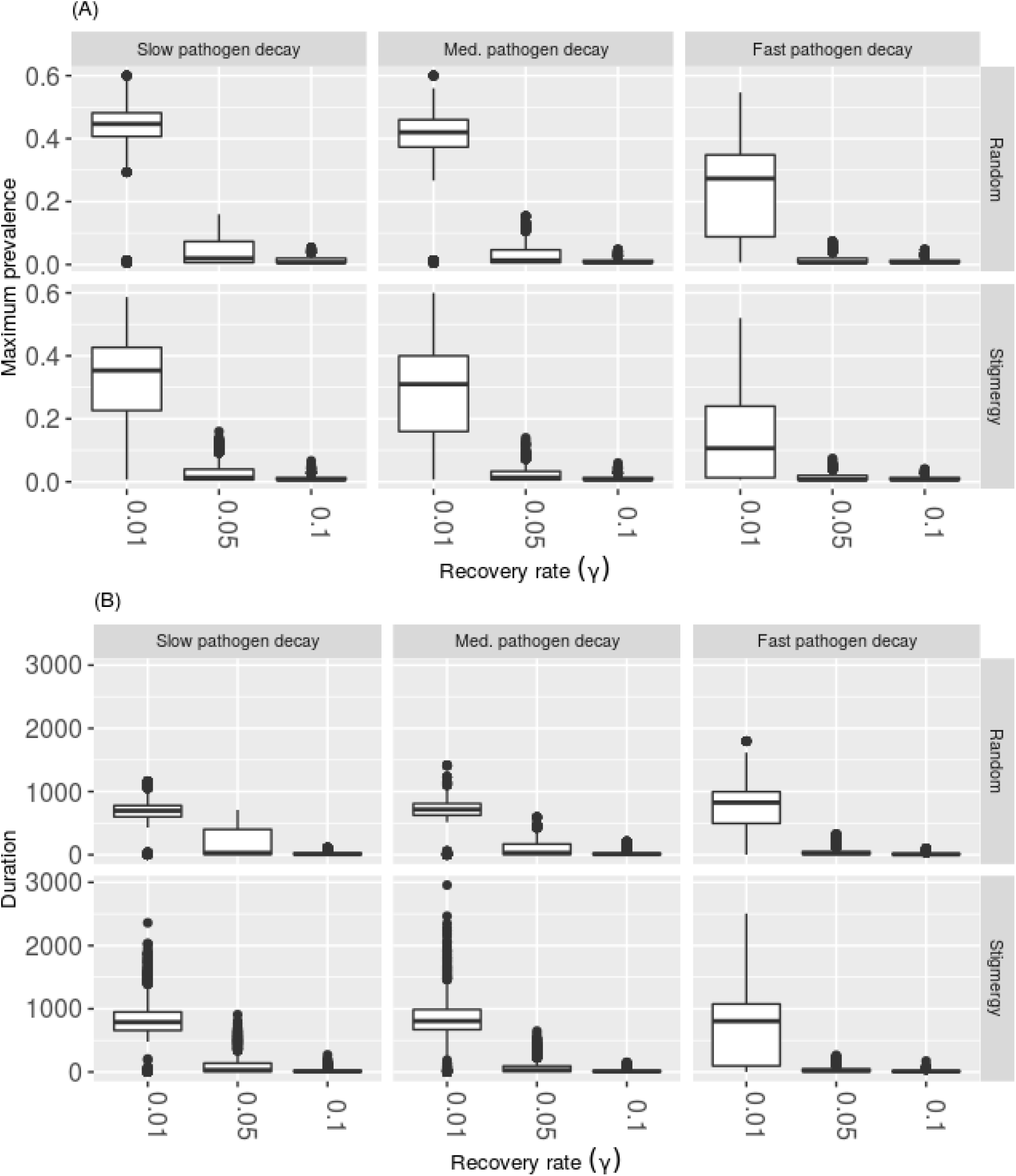
Interaction between environmental decay rate of pathogen (*α*, columns) and stigmergy vs. random movement (*m*, rows) as a function of recovery rate for: (A) maximum prevalence and (B) outbreak duration. Shown for a simulated population size of 150 individuals.

### Territoriality can reduce outbreak severity, but increase disease persistence

Territorial movement yielded lower maximum prevalences in scenarios that were already conducive to outbreaks: a higher density of hosts and slower recovery rates (i.e., longer infectious periods) (Fig 4, S1 Fig). These mitigating effects were strongest for pathogens with faster environmental decay rates prevalence (Fig 4A, S1 Fig). In contrast, at the highest conspecific density treatment, stigmergy increased the variance of outbreak duration, most notably for simulations with slower pathogen decay rates (Fig 4B). These differences in outbreak dynamics between territorial and random movement diminished at lower host densities (S2 Fig).

### Non-linear interactions between pathogen load, pathogen decay, scent load, and scent decay

In the parameter space where outbreaks were most successful (e.g., slower recovery rates and larger population sizes), non-linear patterns emerged from interactions between decay rate of pathogen infectiousness, decay rate of scent cue, initial pathogen load, and initial strength of scent cue. For high conspecific densities (N=100, 150) and the slowest recovery rate (*γ*=0.01), outbreaks reached a higher maximum prevalence for simulations with higher initial pathogen loads, slower pathogen decay rates, lower initial scent loads, and faster scent decay rates (Fig 5A, S3 Fig, lower left quadrant). In contrast, outbreaks lasted longer on average for simulations with higher initial pathogen loads, slower pathogen decay rates, but higher initial scent loads, and slower scent decay rates (Fig 5B, S3 Fig, upper left quadrant). These trends weakened with lower host density and slower recovery rate (S4 Fig). However, at higher host densities with intermediate recovery rates (*γ*=0.05), slow pathogen decay, fast scent decay, and high initial pathogen and scent loads favored longer outbreaks (S5 and S6 Figs). These patterns dissolved for faster recovery rates and lower host densities where outbreaks were less successful (S7–S10 Figs).

**Fig 5.**
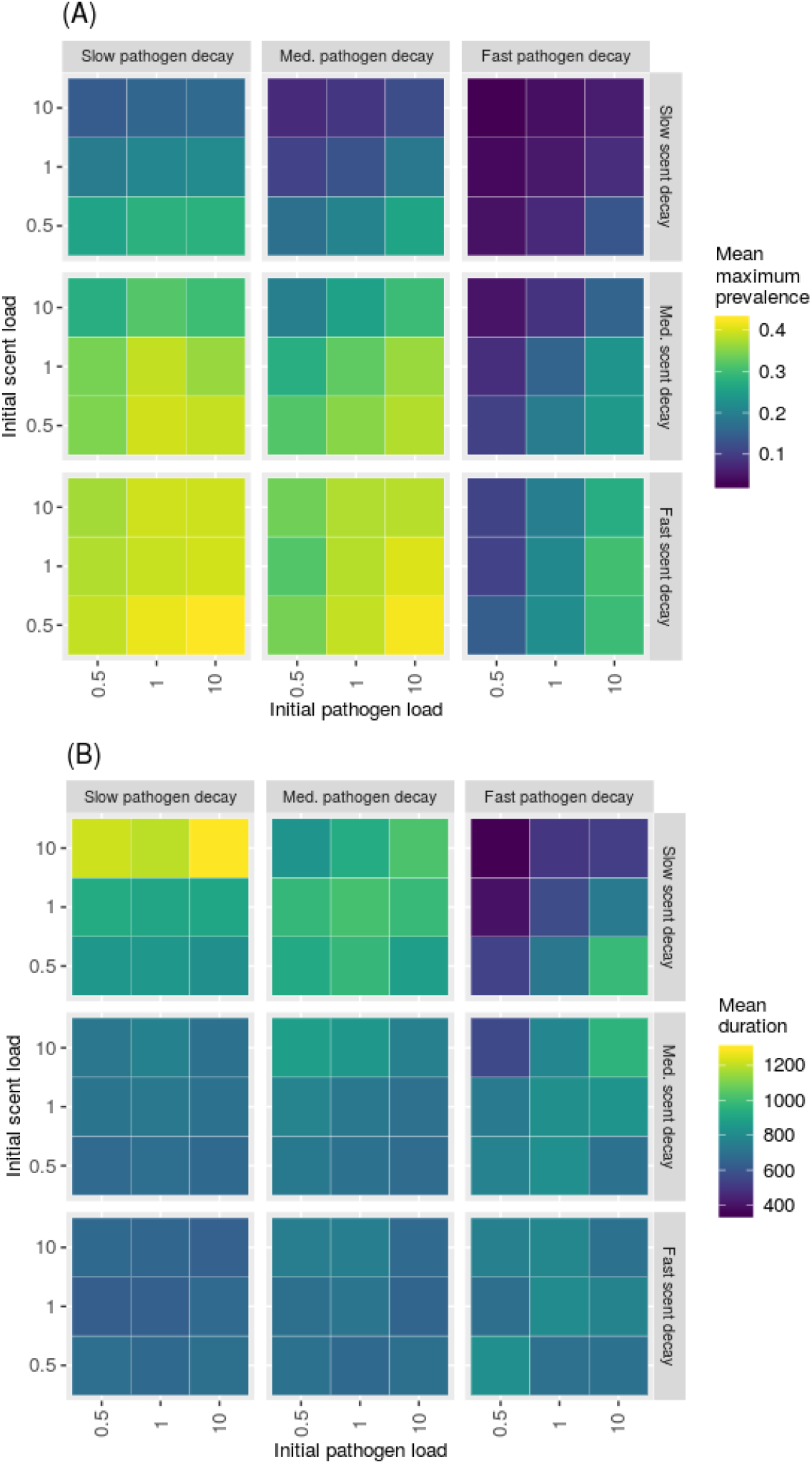
Mean maximum prevalence (A) and mean duration (B) of simulated outbreaks for simulations with 150 individuals responding to stigmergy cues with a recovery rate of 0.01/unit time.

For simulations with slower recovery rate and higher conspecific density, response to initial scent load and pathogen load was variable and interacted with scent and pathogen decay rates (Fig 6, S11 Fig). Together, fast pathogen decay and fast scent load decay rates are not conducive to outbreaks regardless of initial pathogen load or scent load (Fig 6, S11 Fig). Fast scent decay and slow pathogen decay also minimized the effect of different initial pathogen and scent loads (Fig 6, S11 Fig). Lower conspecific density treatments increased variability across outcomes and minimized the differences across scent decay rate, initial pathogen load, and initial scent load for a given pathogen decay rate (S12 Fig).

**Fig 6.**
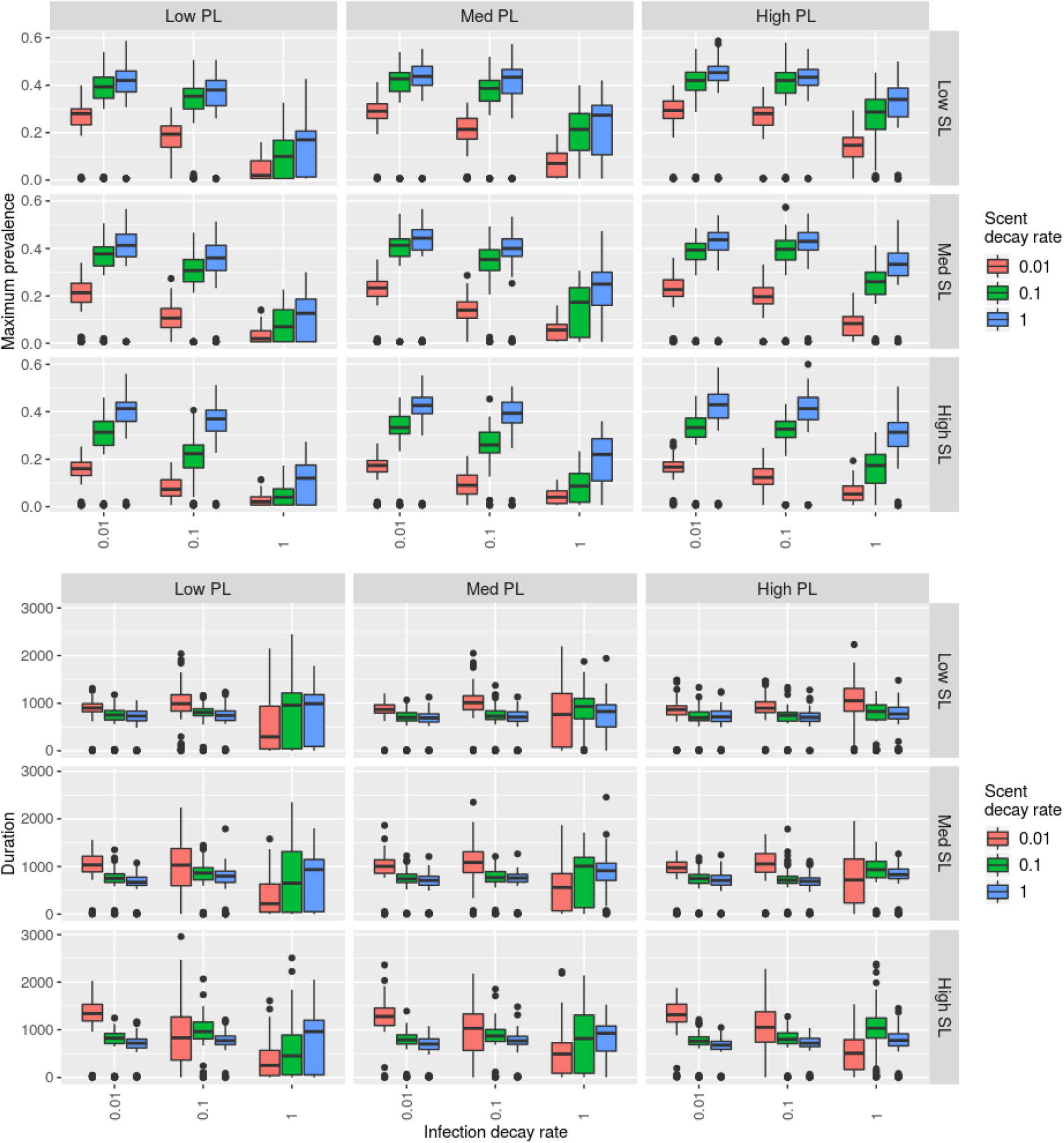
Boxplots of (A) maximum prevalence and (B) outbreak duration with 150 simulated individuals responding to stigmergy cues and a recovery rate of 0.01/time step. Rows correspond to low, medium, and fast scent loads (SL). Columns correspond to low, medium, and fast pathogen loads (PL).

## Discussion

Adaptive, dynamic, or territorial space use is one possible explanation for the lack of empirically observed density thresholds in wildlife [26, 27]. This “territoriality benefits” hypothesis suggests that reduced home range overlap could lead to reduced opportunities for pathogen transmission [32]. Our findings support this hypothesis with the caveat that a reduction in outbreak severity may come at the cost of increased likelihood of persistence for indirectly transmitted pathogens. We found that territoriality did indeed reduce maximum prevalence of disease in conditions where we would otherwise expect outbreaks to be most successful: slower recovery rates (i.e., longer infectious periods) and higher conspecific densities (Fig. 4, S1 Fig). However, at high enough densities, outbreak duration becomes considerably more variable for populations with stigmergy-driven movement compared to their randomly moving counterparts (Fig. 4).

For longer infectious periods and higher host densities, key trade-offs emerged between the strength of pathogen load, strength of the stigmergy cue, and the rate at which those two quantities decayed. Intuitively, high initial pathogen load and a slower pathogen decay rate universally promoted higher maximum prevalence and longer lasting outbreaks (Fig. 5 & 6). In contrast, higher initial scent loads paired with faster scent decay promoted higher maximum prevalences (Fig. 5A), whereas higher initial scent loads and slower scent decay rates promoted longer lasting outbreaks (Fig. 5B). These findings raise interesting questions about the evolutionary nature of the competing strength of pathogen and scent marking signals in the environment. Indirectly transmitted pathogens should coevolve for longer persistence and higher virulence because individual host mortality is less important to a pathogen’s overall fitness [33]. This is in opposition to the prediction that populations with spatially restricted movement will contribute to the evolution of less virulent pathogens [33]. Our results support the idea that pathogens co-opting their hosts’ social communication system could help to overcome territorial barriers (e.g., [29]) and that territorial behavior could offer benefits for stochastic persistence from a pathogen’s evolutionary perspective (Fig. 4B). In particular, a host’s tendency to deposit less strong, but more slowly decaying scent mark could help maintain pathogen persistence (Fig 5B).

In our model, indirectly transmitted pathogen dynamics could be altered by territorial cues if hosts existed at high enough densities and shed pathogens for long enough across the landscape. This work, therefore, highlights the importance of exploring feedbacks between territoriality and parasitism. In empirical systems, hosts with lower levels of parasitism may be better able to form and maintain territories. For example, pheasants are a competent host for Lyme disease and commonly parasitized by *Ixodes ricinus* ticks. Male pheasants with experimentally reduced tick loads were more likely to gain harems and have smaller territories. In contrast, males with higher tick loads ranged more broadly in peripheral woods and fields leading to a positive feedback loop of higher likelihood of tick exposure [34]. Examples of negative feedback between parasitism and territoriality also exist. In male Grant’s gazelle (*Nanger granti*), territorial behavior drives higher parasite loads, but higher parasite loads suppress behaviors associated with territoriality [35]. Future model development might consider incorporating such feedback mechanisms [36], e.g., differences in movement behavior between symptomatic and healthy individuals.

To highlight the competing axes of stigmergy cue strength and duration vs. pathogen load strength and duration, we simulated movement using a random walk rather than incorporating additional potential complexities of movement behavior; this necessarily means that simulated individuals did not respond to the real-time presence or absence of conspecifics in neighboring cells. Future modelling studies could explore the sensitivity of results to differences in perceptual range (i.e., extending beyond a Moore neighborhood) and memory of past movements or past stigmergy cue encounters. Other extensions might include accounting for dispersal behavior or inter-individual differences in home range size. Ultimately, stigmergy is just one possible mechanism for informing territorial-like movement behavior. It is likely that many species respond to cues in real time (e.g., visual cues, vocalization) in addition to transient environmental cues (e.g., [37]). Another important question is understanding how temporal switches in the valence of the stigmergy cues might affect disease transmission. For example, during mating seasons scent cues could become attractive rather than aversive [20]. Individuals are also likely to display heterogeneous responses to different members of the population (e.g., male vs. female) and their environmental cues [38].

The model presented here best describes indirect or environmental transmission of a single infectious agent within an asocial, territorial host species. However, this model could also describe the behavior of social territorial carnivores (e.g., gray wolves, African lions), where the movement of a single individual is usually representative of the entire group [39]. This model framework may also be relevant for pathogens with other dominant transmission modes that persist in the environment for extended periods. For example, canine parvovirus, which can persist up to one year outside of a host [40], and is of conservation concern for wild carnivores [41]. Similarly, leptospirosis, a bacterial infection of wildlife (and humans), can persist for months in aqueous environments [42]. Small mammals, including peri-domestic species like raccoons, secrete bacteria through urine [43, 44], which can serve as a scent marking communication tool [45]. Likewise, feline calicivirus remains infectious for up to 20 days at ambient temperatures [46] and is of epidemic concern for African lions [18]. Some domestic cats remain persistently infected (shedding virus for more than 30 days) and may shed higher levels of virus [16], which corresponds to the slower recovery rate condition in our model.

In an applied context, scent marking behavior can serve as a way to assess animal populations through time and document responses to human disturbance [29, 47]. Our results support the idea that decision makers should evaluate possible changes in scent marking behavior and its potential effects on disease control when considering culling or altering population size in territorial species [48, 49]. For example, prior attempts to control bovine tuberculosis (*Mycobacterium bovis*) in badgers through culling caused changes in scent marking behavior. At lower densities, badgers were more likely to have dispersed patterns of fecal and urine scent marking with higher concomitant risks of pathogen transmission [50]. Bovine tuberculosis transmission is thought to occur primarily through direct contact, but its ability to persist in the environment has raised questions about the role of indirect transmission routes [51, 52]. Likewise, wildlife scent marking behavior at the human-livestock interface is of concern since some wildlife species (e.g., foxes and badgers) preferentially use farm food storage buildings for foraging and scent-marking which heightens pathogen transmission risk [53,54]. Scent marking may also influence the success of species reintroductions and population management: introducing translocated animals into an established territorial population may increase transmission risk because of increased overlap in home ranges or direct contacts [55]. Similarly, anthropogenic resource supplementation increases risk of indirect or fomite transmission (e.g., bovine tuberculosis in deer, brucellosis in elk, chronic wasting disease in deer and elk) [56]. An interesting question moving forward would be to investigate the competing roles of habitat quality and territoriality on disease dynamics for pathogens with environmental persistence [57].

Existing movement ecology studies have so far focused on how to model territorial behavior and not the consequences of dynamic territories on population-level outcomes like disease [2]. Mechanistic portrayals of how animals form and maintain territories remain rare, and to our knowledge, have not been integrated with a disease modeling framework. This work provides a key interface between the disciplines of movement and disease ecology [4, 5, 13] by exploring how mechanistic movement driven by an individual’s social landscape affects disease dynamics. These results indicate an interesting threshold at higher host densities where stigmergy-driven movement behavior can still support pathogen persistence. This framework can be adapted to specific host–pathogen systems to generate hypotheses about the competing roles of transient social cues and indirect pathogen transmission. We hope that this model inspires additional research surrounding the role of socially driven movement behavior and its concomitant implications for pathogen transmission.

## Methods

### Initial conditions and parameter space

We simulated a 50 x 50 cell landscape with wrapped edges (i.e., torus) to avoid boundary effects [58]. At the start of each simulation, individuals were randomly distributed across the theoretical landscape, and one randomly infected individual served as the index case. We tested population sizes of 50, 100, and 150 individuals for conspecific densities of 0.02, 0.04, and 0.06 hosts/unit area respectively. Since the strength of the encountered pathogen load controlled the transmission probability, we explored the epidemiological parameter space by simulating low, medium, and high recovery rates (Table 1). We also explored the interplay of low, medium, and high deposition strengths for scent marking and pathogen shedding, as well as low, medium, and high rates of decay for pathogen infectiousness and scent mark strength (Table 1).

### Movement

At each time step, individuals could move within a Moore neighborhood (8 neighboring cells) or remain within their current cell. Movement was weighted by the simplest case of a movement kernel: *ϕ*(*r*) = 1/(2π*r*^2^) where the weight, *ϕ*(*r*), is inversely proportional to the radial distance (*r*) from the center point of the current grid cell: 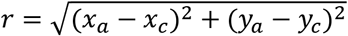

The equation gives an inverse distance weight (i.e., 1/*r*) that is multiplied by the circumference at that distance to account for a uniform circular distribution (i.e., 1/(2*πr*)) [57]. By setting the minimum distance (*r*)for the cell of origin to 0.75, hosts were slightly more likely to to remain in the current cell (p=0.23) than move in one of the four cardinal directions (p=0.13) or move in a diagonal direction (p=0.06).

### Variable importance analysis

We explored model sensitivity to parameter values by conducting a random forest variable importance analysis. Random forest analysis is an approach that accounts for non-linear and collinear relationships between variables, allows for different variable types (e.g., numerical vs. categorical), and avoids the concerns of using frequency-based statistical p-values to assign significance in a simulation context [59,60]. Here we used the *party* package, which has been shown to be particularly robust to bias relative to the traditional *randomforest* package [61–63]. Results are reported as mean decrease in accuracy scores, which describes the loss in accuracy to the predicted outcome when the given variable is permutated randomly [59]. We used the *cforest* function and the conditional *varimp* function for generation of 1,000 trees for the metrics of outbreak success, maximum prevalence, and duration. With 1,000 trees, the order of variable importance did not switch with different random seeds. All simulations and analyses were conducted in R (version 3.5.3). Code is available at: https://github.com/whit1951/StigmergyDisease and will be uploaded to Zenodo upon acceptance.

---

**S1 Fig.**
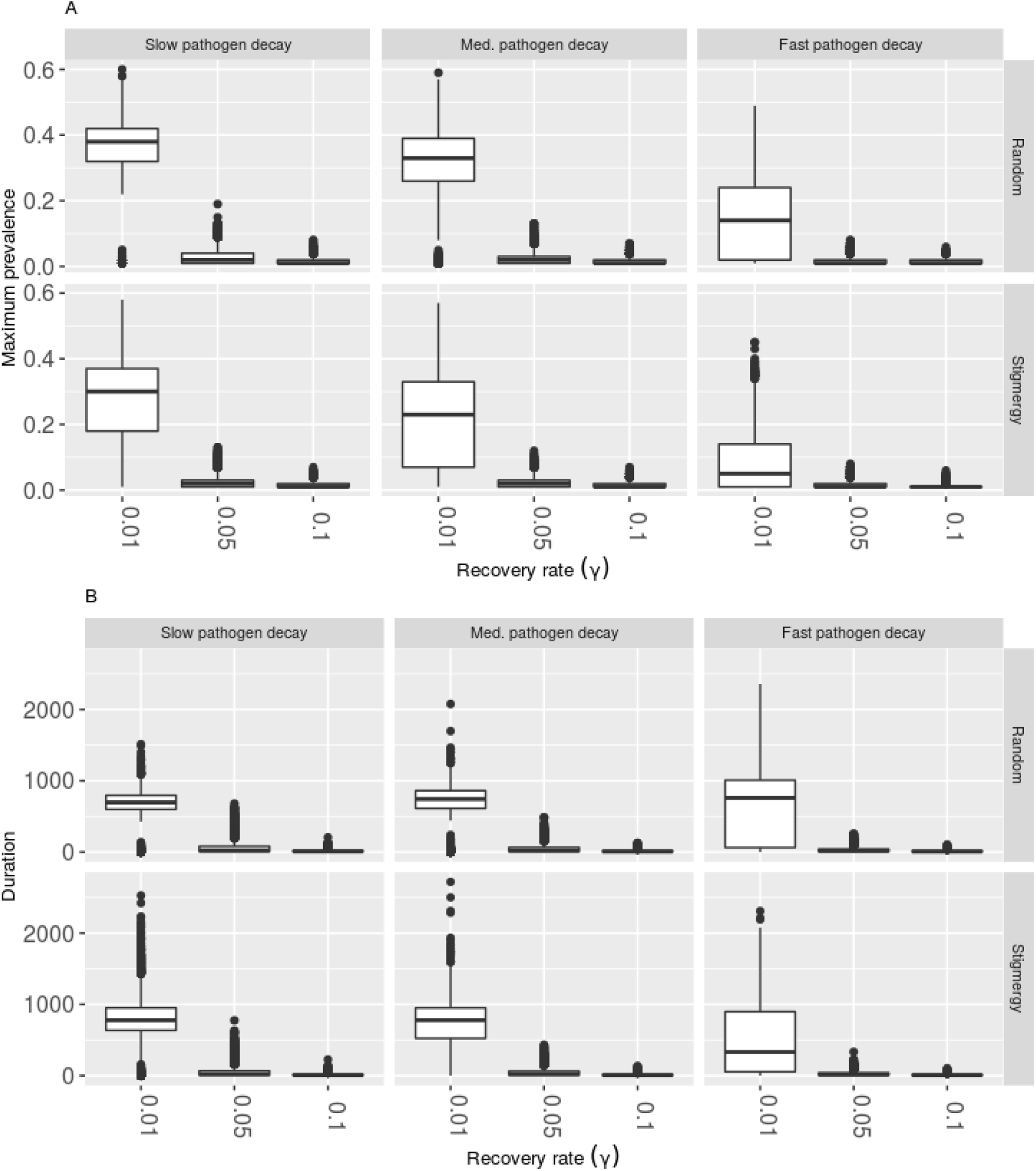
Interaction between environmental decay rate of pathogen (*α*, columns) and stigmergy vs. random movement (*m*, rows) as a function of recovery rate for (A) maximum prevalence and (B) outbreak duration. Shown for a simulated population size of 100 individuals.

**S2 Fig.**
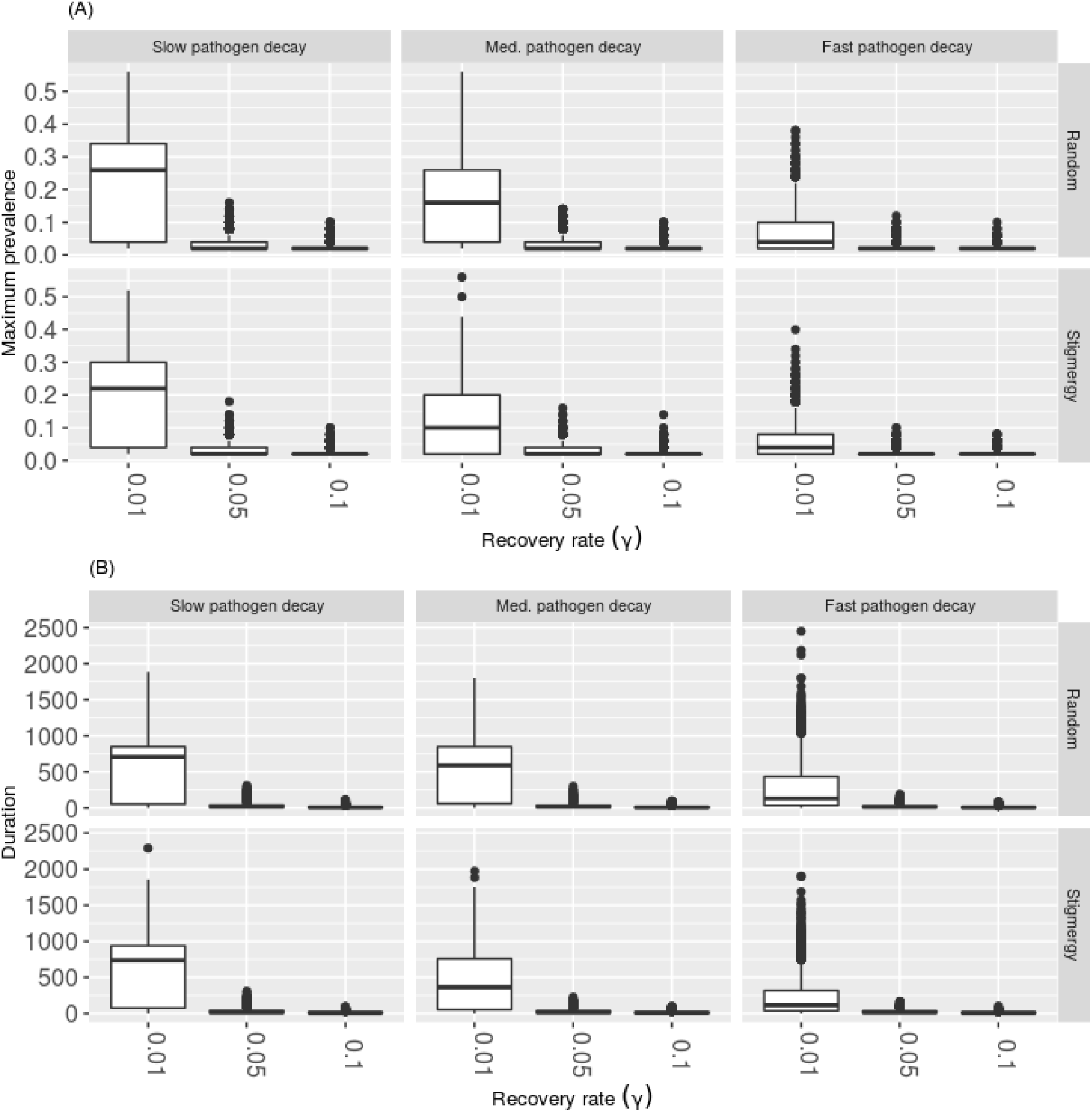
Interaction between environmental decay rate of pathogen (*α*, columns) and stigmergy vs. random movement (m, rows) as a function of recovery rate for (A) maximum prevalence and (B) outbreak duration. Shown for a simulated population size of 50 individuals.

**S3 Fig.**
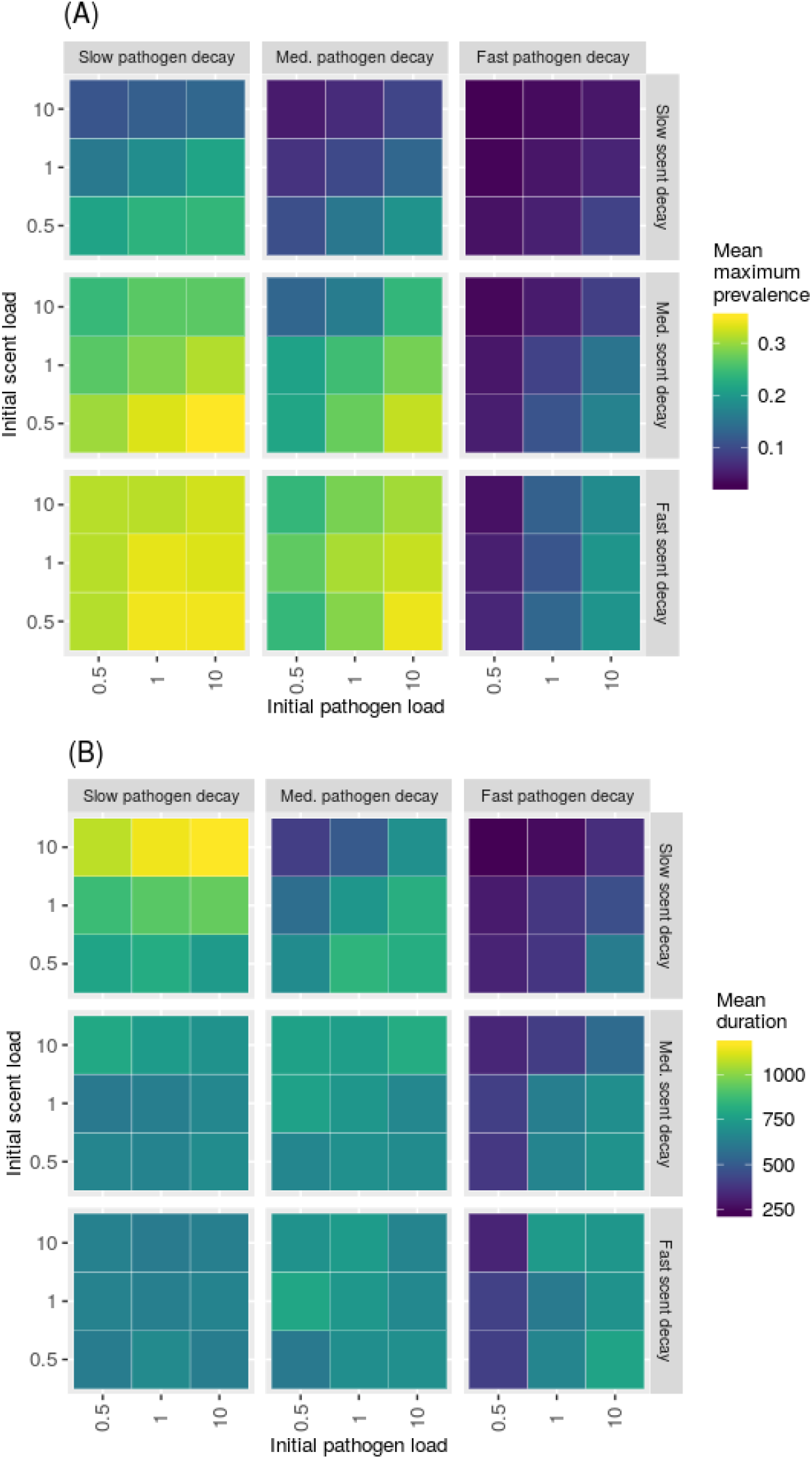
Mean maximum prevalence (A) and mean duration (B) of simulated outbreaks for simulations with 100 individuals responding to stigmergy cues with a recovery rate of 0.01/unit time.

**S4 Fig.**
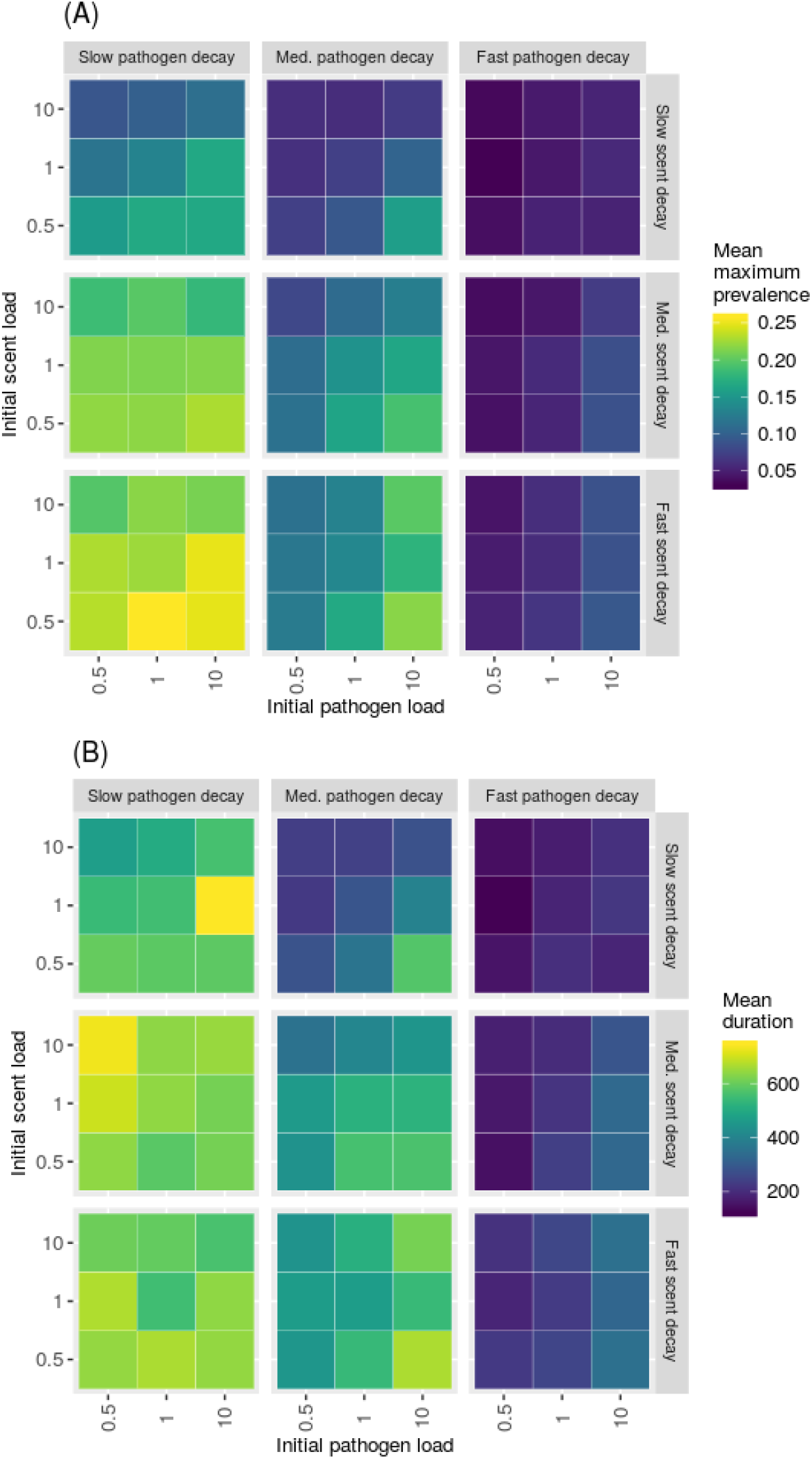
Mean maximum prevalence (A) and mean duration (B) of simulated outbreaks for simulations with 50 individuals responding to stigmergy cues with a recovery rate of 0.01/unit time.

**S5 Fig.**
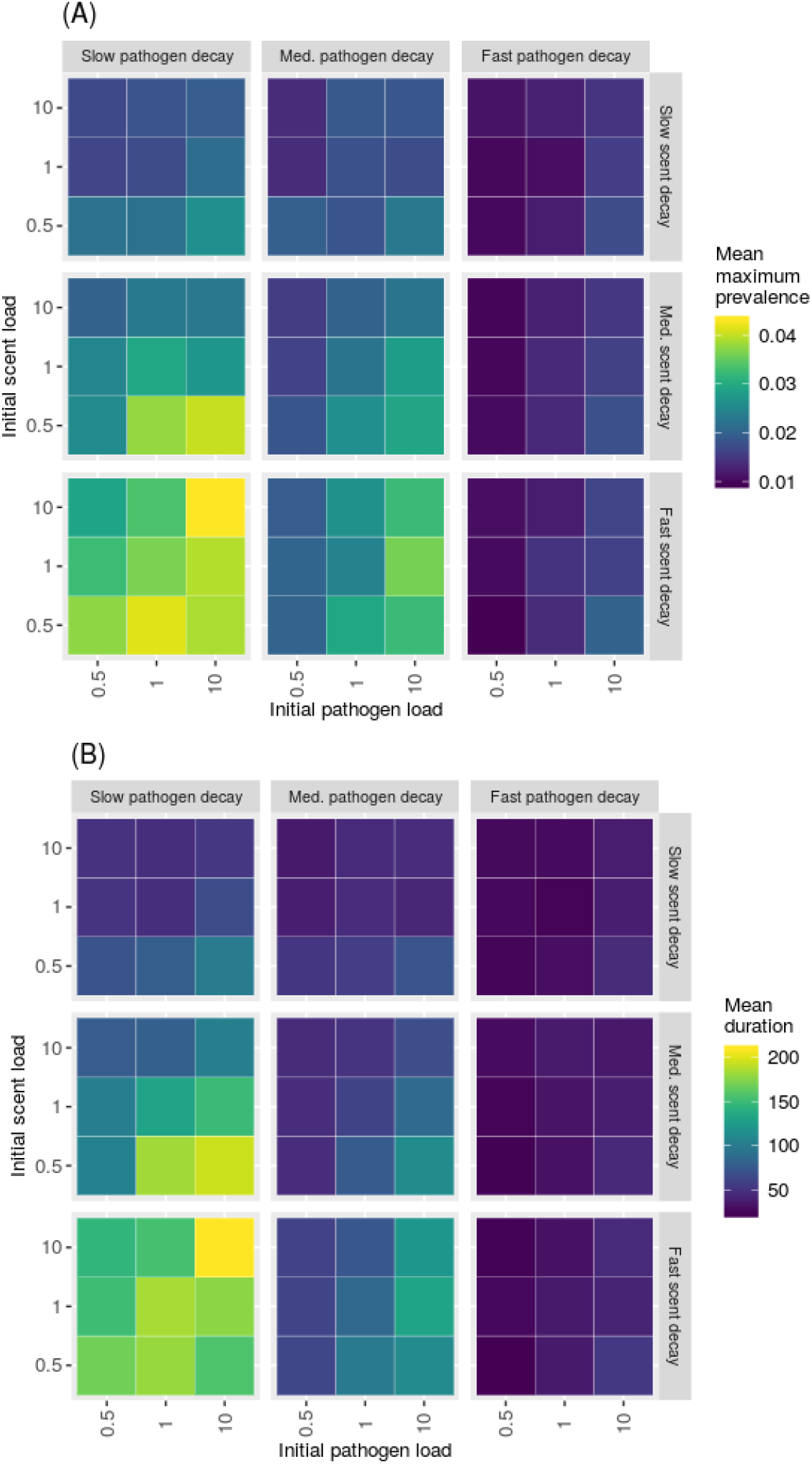
Mean maximum prevalence (A) and mean duration (B) of simulated outbreaks for simulations with 150 individuals responding to stigmergy cues with a recovery rate of 0.05/unit time.

**S6 Fig.**
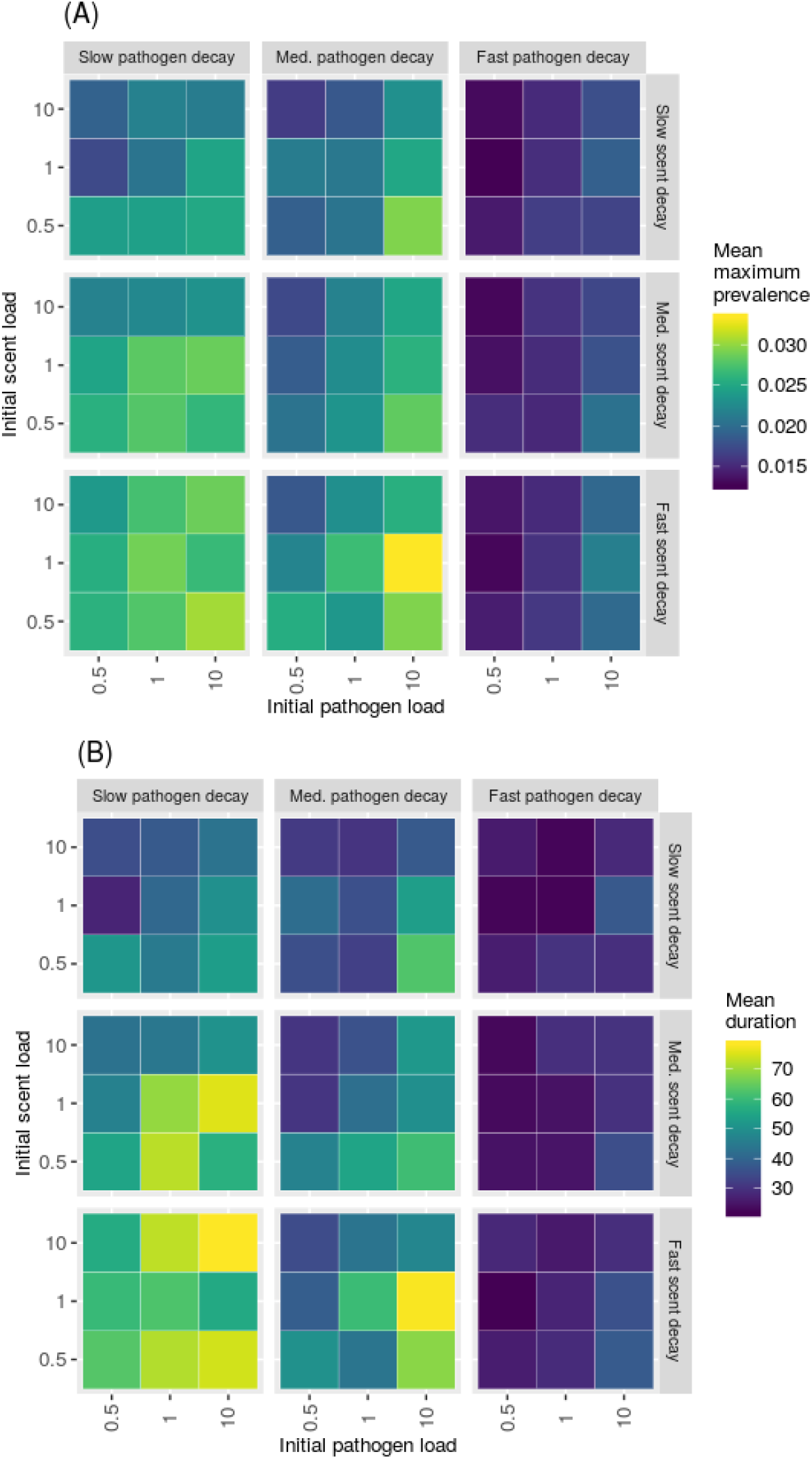
Mean maximum prevalence (A) and mean duration (B) of simulated outbreaks for simulations with 100 individuals responding to stigmergy cues with a recovery rate of 0.05/unit time.

**S7 Fig.**
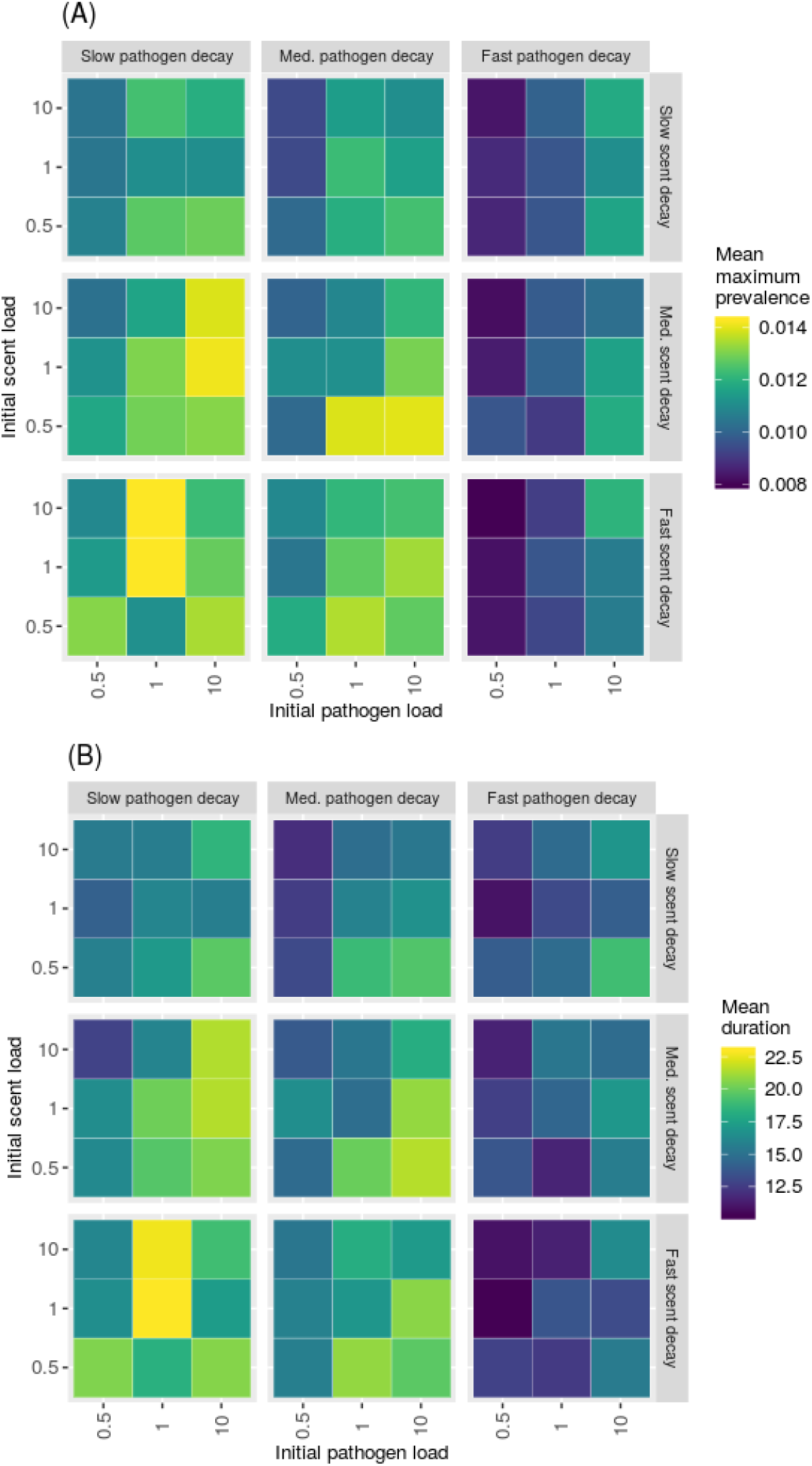
Mean maximum prevalence (A) and mean duration (B) of simulated outbreaks for simulations with 150 individuals responding to stigmergy cues with a recovery rate of 0.10/unit time.

**S8 Fig.**
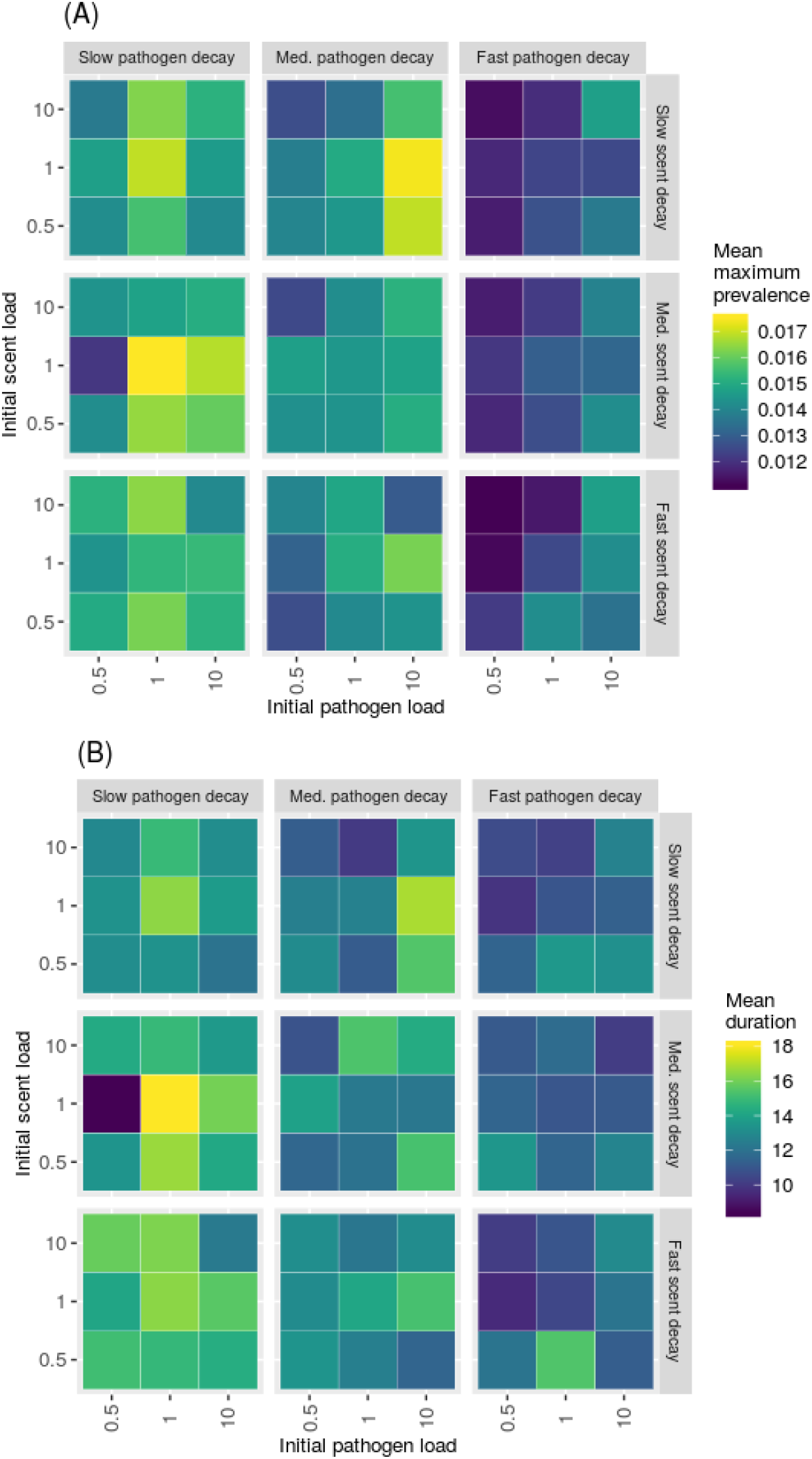
Fig. Mean maximum prevalence (A) and mean duration (B) of simulated outbreaks for simulations with 100 individuals responding to stigmergy cues with a recovery rate of 0.10/unit time.

**S9 Fig.**
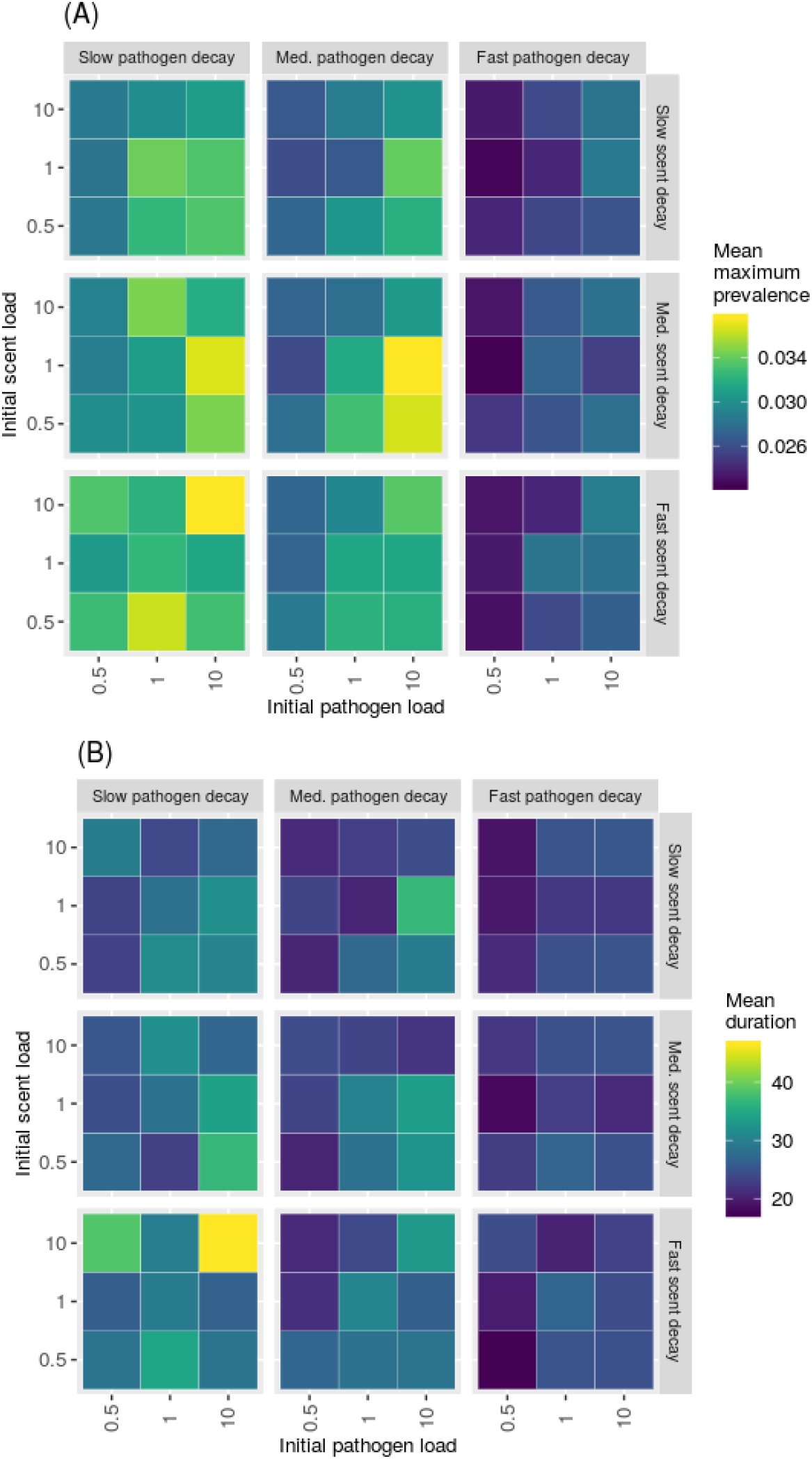
Mean maximum prevalence (A) and mean duration (B) of simulated outbreaks for simulations with 50 individuals responding to stigmergy cues with a recovery rate of 0.05/unit time.

**S10 Fig.**
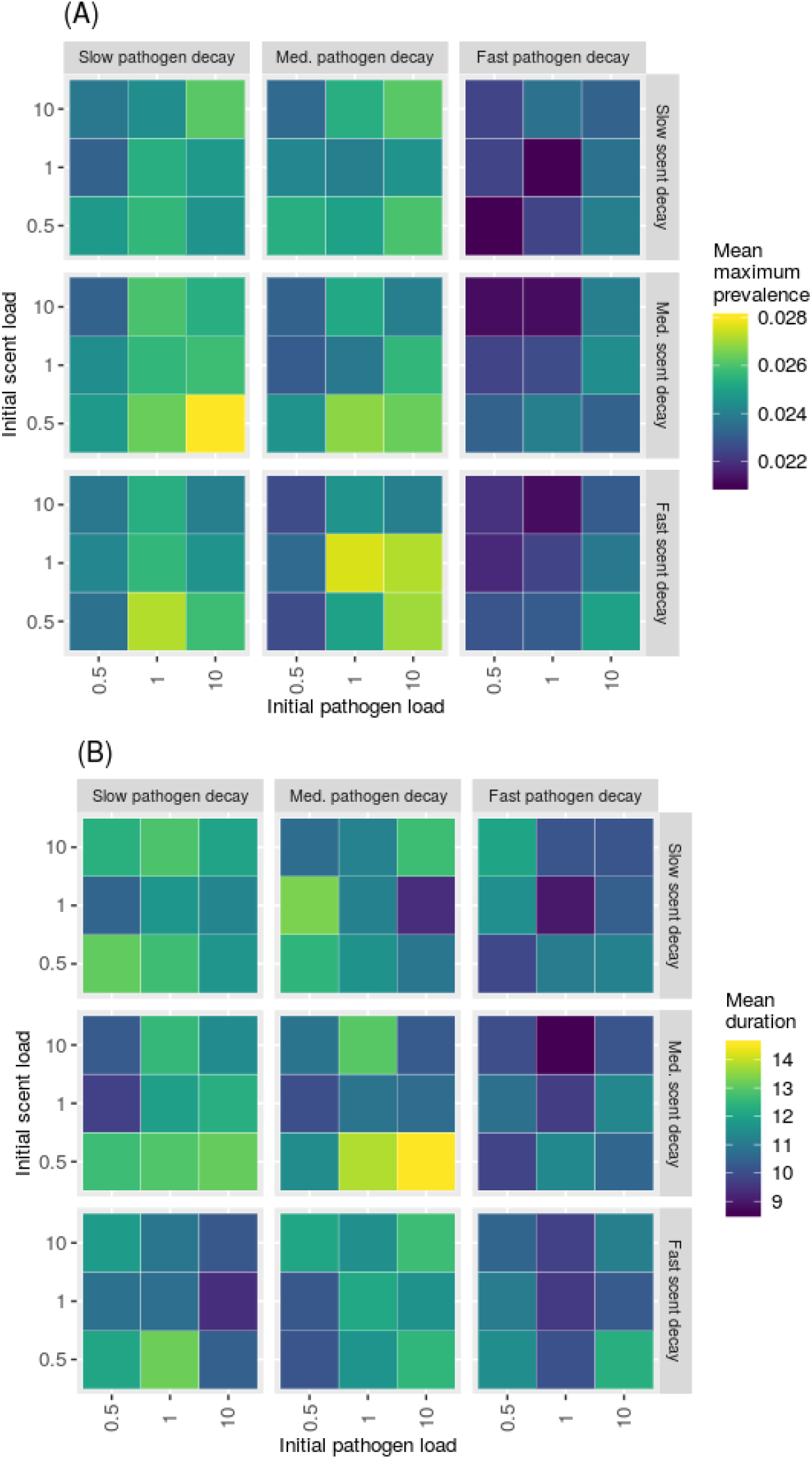
Mean maximum prevalence (A) and mean duration (B) of simulated outbreaks for simulations with 50 individuals responding to stigmergy cues with a recovery rate of 0.10/unit time.

**S11 Fig.**
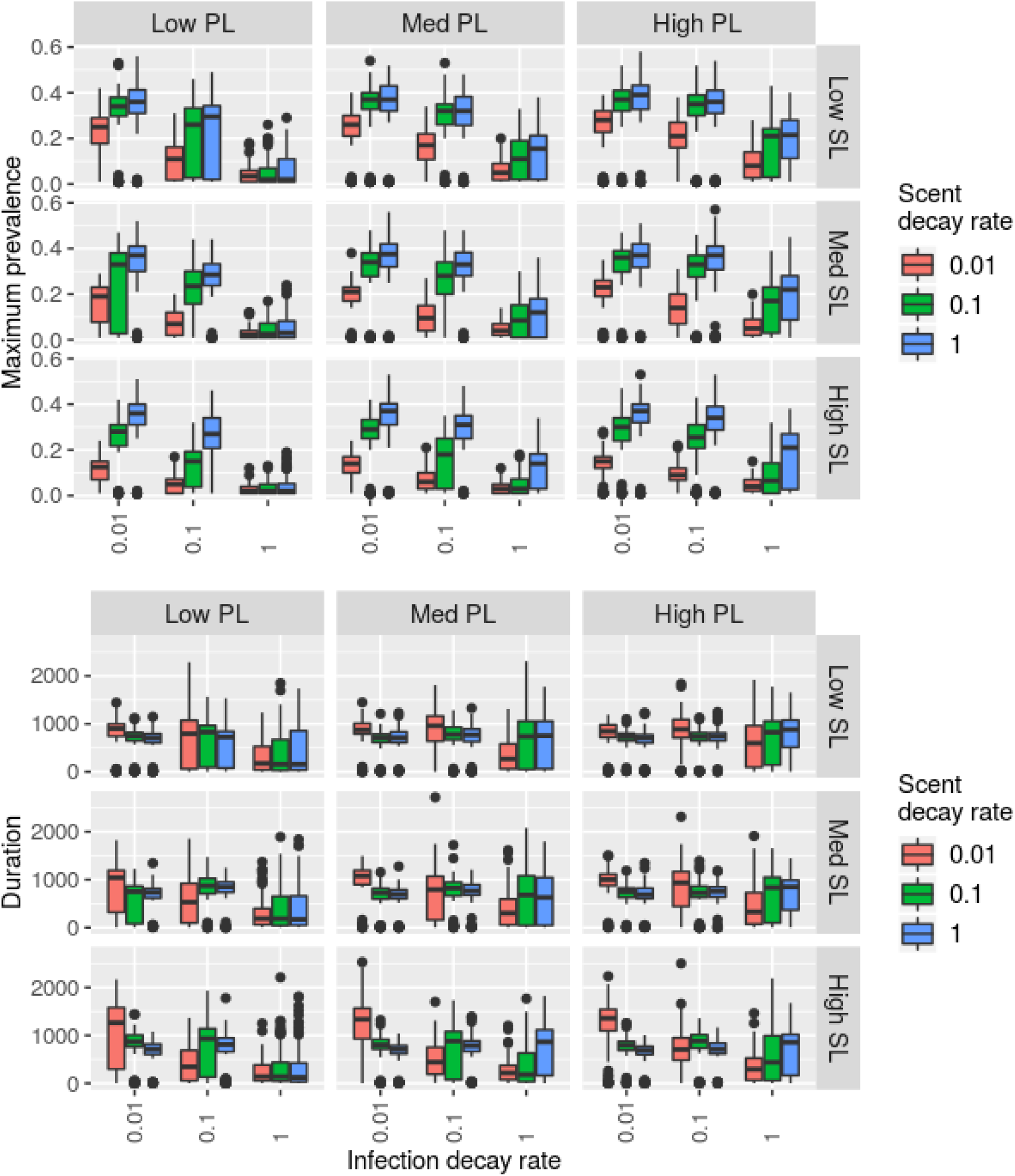
Boxplots of (A) maximum prevalence and (B) outbreak duration with 100 simulated individuals responding to stigmergy cues and a recovery rate of 0.01/time step. Rows correspond to low, medium, and fast scent loads (SL). Columns correspond to low, medium, and fast pathogen loads

**S12 Fig.**
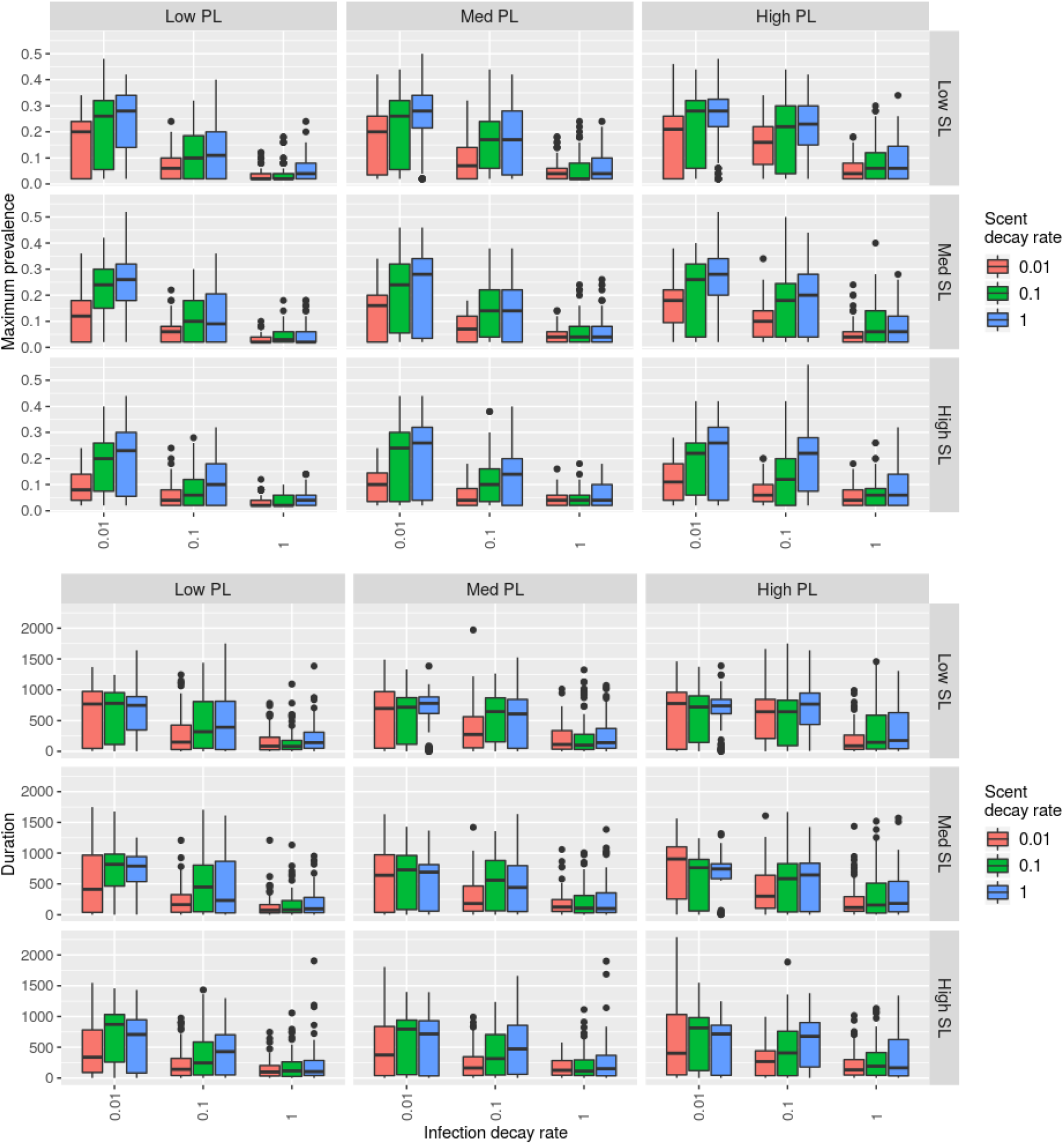
Boxplots of (A) maximum prevalence and (B) outbreak duration with 50 simulated individuals responding to stigmergy cues and a recovery rate of 0.01/time step. Rows correspond to low, medium, and fast scent loads (SL). Columns correspond to low, medium, and fast pathogen loads

